# Myristoylation alone is sufficient for PKA catalytic subunits to fractionally associate with the plasma membrane to regulate neuronal functions

**DOI:** 10.1101/2020.06.25.172676

**Authors:** Wei-Hong Xiong, Maozhen Qin, Haining Zhong

## Abstract

Myristoylation is a post-translational modification that plays diverse functional roles in many protein species. The myristate moiety is considered insufficient for protein-membrane associations unless additional membrane-affinity motifs, such as a stretch of positively charged residues, are present. Here, we report that the electrically neutral N-terminal fragment of the protein kinase A catalytic subunit (PKA-C), in which myristoylation is the only functional motif, is sufficient for membrane association. This myristoylation can associate a fraction of PKA-C molecules or fluorescent proteins (FPs) to the plasma membrane in neuronal dendrites. The net neutral charge of PKA-C is evolutionally conserved, even though its membrane affinity can be readily tuned by changing charges near the myristoylation site. The observed membrane association, while moderate, is sufficient to concentrate PKA activity at the membrane by nearly 20-fold, and is required for PKA regulation of AMPA receptors at neuronal synapses. Our results indicate that myristoylation alone may be sufficient to drive functionally significant membrane association in the absence of assisting motifs. This provides a revised foundation for the understanding of how myristoylation regulates protein functions.

## Introduction

Myristoylation is a major type of post-translational modification that occurs at the N-terminus of a myriad of proteins (Carr et al., 1982; Farazi et al., 2001; Resh, 2013; Wright et al., 2010). Depending on the target, myristoylation can contribute to the structure, stability, protein-protein interactions, and subcellular localization of the modified proteins (Farazi et al., 2001; Resh, 2013; Wright et al., 2010). In particular, myristoylation often facilitates protein association with the membrane. However, it is thought that, with an acyl chain of only 14 carbons, myristate confers insufficient energy for stable association of a protein with the membrane (Murray et al., 1997; Peitzsch and McLaughlin, 1993). Subsequent studies have shown that a second membrane-affinity motifs, such as a stretch of basic residues or a second lipid modification, is required for the membrane association of several myristoylated proteins (reviewed in Farazi et al., 2001; Resh, 2013; Wright et al., 2010). When the second membrane-affinity motif is removed or neutralized, either physiologically or via mutagenesis, the membrane localization of the protein is disrupted. Thus, the canonical view is that myristoylation alone is not sufficient to provide a functionally significant association of a protein with the plasma membrane, even though myristoylation has been observed to be associated with reconstituted lipid bilayers (Struppe et al., 1998).

Myristoylation was first discovered in the catalytic subunit of protein kinase A (PKA) (Carr et al., 1982; Francis and Corbin, 1994), which is a primary mediator of the second messenger cAMP that plays diverse essential roles in nearly all organisms, from bacteria to humans. At rest, PKA is a tetrameric protein that consists of two regulatory subunits (PKA-R) and two catalytic subunits (PKA-C). PKA holoenzymes are anchored to specific subcellular locations via the binding of PKA regulatory subunits (PKA-Rs) with A-Kinase anchoring proteins (AKAPs) (Lohmann et al., 1984; Scott and Pawson, 2009; Theurkauf and Vallee, 1982; Wong and Scott, 2004). In the presence of cAMP, PKA-C is released from PKA-R and becomes an active kinase (Beavo et al., 1974; Francis and Corbin, 1994; Johnson et al., 2001; Naira et al., 1985; Tillo et al., 2017).

Despite being myristoylated, PKA-C is thought to function as a cytosolic protein because of its high solubility (Johnson et al., 2001; Naira et al., 1985). Consistently, a PKA-C mutant with disrupted myristoylation has been shown to perform catalytic function and to maintain several PKA functions in heterologous cells (Clegg et al., 1989). Structural studies found that PKA-C myristoylation site is folded into a hydrophobic pocket, likely to serve a structural role (Bastidas et al., 2012; Yonemoto et al., 1993; Zheng et al., 1993). This view has started to shift based on recent reports showing that activated PKA-C can associated with the membrane in a myristoylation-dependent manner (Gaffarogullari et al., 2011; Tillo et al., 2017; Zhang et al., 2015), including in neurons. However, the extent to which PKA-C associates with the plasma membrane in living cells and its functional significance are not known. Furthermore, as discussed above, myristolyation-mediated membrane association is thought to require a second membrane motif. The identity of this second membrane-affinity motif has not been determined. Therefore, we set out to address these questions.

## Results

First, we found that the N-terminal 47 residues of PKA-C (named PKA-Cn) (Fig. 1a, upper panel), which lacks the kinase domain and the C-terminal domains and cannot bind to PKA-R, was sufficient to target an FP to the plasma membrane in neuronal dendrites. Specifically, cultured hippocampal slices were transfected with PKA-Cn fused to the N terminus of yellow FP mVenus (PKA-Cn-mVenus) together with a red cytosolic marker (mCherry). The thick (ϕ >= 1.5 µm) apical dendrites of CA1 neurons were imaged live at 2– 4 days post-transfection under a two-photon microscope. An optical section through the center of the dendrite (Fig. 1b, left panels) was used to compare the localizations of PKA-Cn-mVenus and mCherry. The PKA-C-mVenus fluorescence (green) was more enriched at the periphery of the dendrite when compared to mCherry fluorescence (red, shown as magenta). To quantify the membrane association, a membrane enrichment index (MEI) was computed:

**Figure 1.**
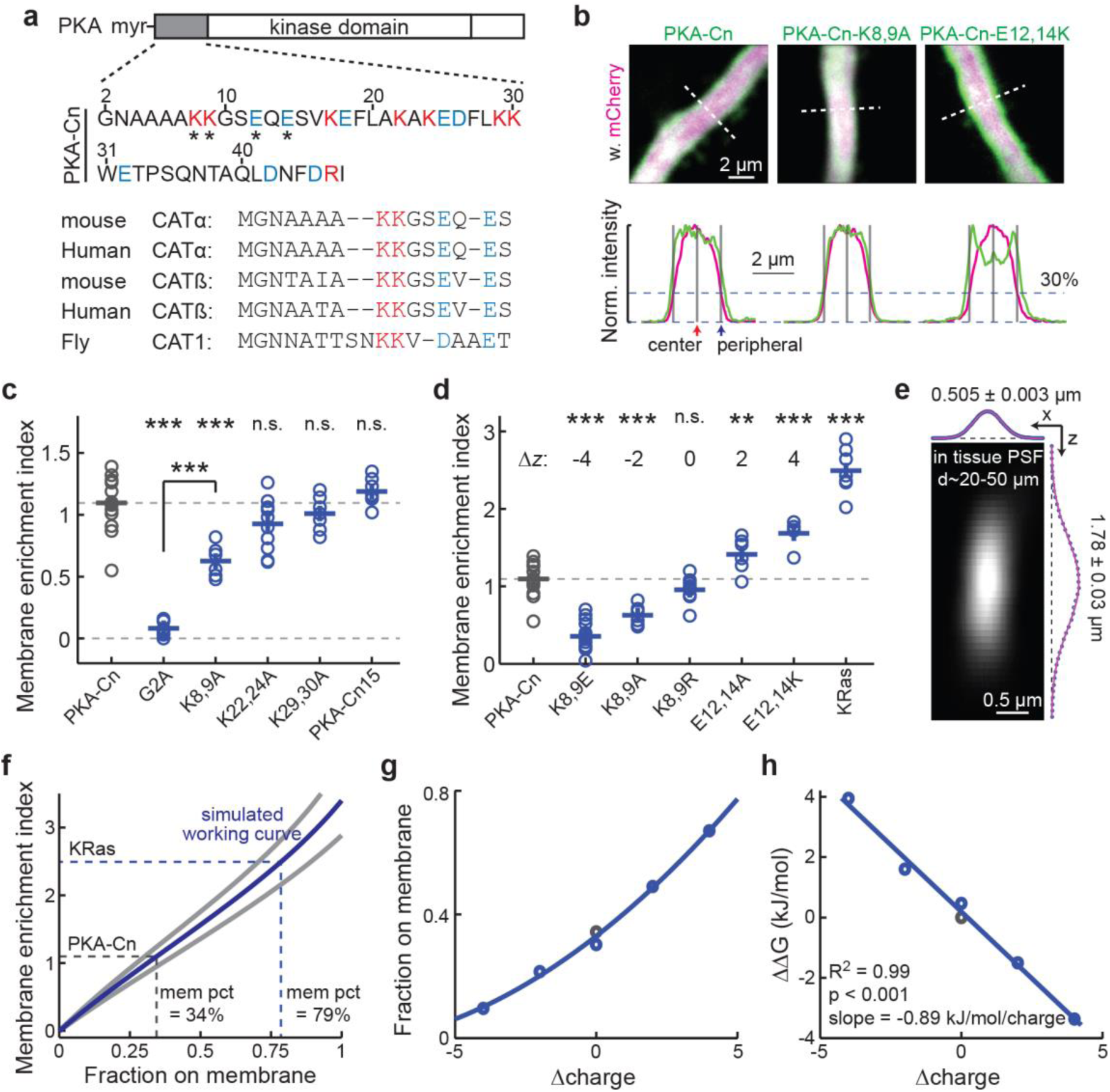
PKA-Cn is sufficient to localize a fraction of fluorescent proteins to the plasma membrane. **a**, Sequence of PKA-C N terminus showing balanced positive (magenta) and negative (blue) charges. Upper: the 47 residues of mouse PKA-C used in this study. The residues affecting membrane affinity are indicated (asterisks). Lower: alignment of the first 15 residues of the indicated PKA-C subtypes and species. **b**, Representative images and traces along the indicated dashed white lines of thick neuronal apical dendrites transfected with the indicated constructs for quantifying MEIs. The red fluorescence of mCherry is shown as magenta (to accommodate color-blind readers). Gray bars mark the locations where the fluorescence intensities are used to calculate MEIs: at the center and at the periphery where the red signal is at 30% of peak (corresponding to the plasma membrane)(Tillo et al., 2017). **c**, Collective MEI quantifications for the indicated PKA-Cn-mVenus constructs. From left to right, n = 16, 12, 6, 11, 7, and 7 for the indicated constructs. **d**, Collective MEI quantifications for the indicated PKA-Cn-mVenus constructs and KRas-mEGFP. The charge relative to PKA-Cn (Δ*z*) is indicated for each construct. The data for PKA-Cn is the same as in panel c. For other constructs, from left to right, n = 18, 6, 11, 6, 5, and 7. **e**, Measured PSF in cultured hippocampal slices at imaging depths of 20-50 µm. n = 36. **f**, MEI versus membrane fraction working curves computed based on simulation using the measured PSF size and the experimental dendritic width of PKA-Cn at mean (blue) and one standard deviation (gray). As indicated, the percentages of PKA-Cn and KRas on the membrane (mem pct) were estimated based on the mean-size curve. **g**, Charge versus membrane fraction relationships for the PKA-Cn constructs in panel d. Membrane fractions were estimated based on the mean MEI values and the corresponding dendrite sizes. **h**, The relationship between the charge versus Gibbs free energy relative to wild-type PKA-Cn for the constructs in panel g, and its fit to a linear function.

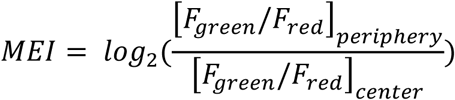

where F is the fluorescence intensity. The peripheral fluorescence intensities were measured at the edge of the dendrite, as determined by the location where the red was 30% of the peak fluorescence (Fig. 1b, lower panels) (Tillo et al., 2017). MEI values greater than 0 would indicate membrane enrichment, whereas MEI values less than 0 would indicate membrane exclusion. PKA-Cn-mVenus exhibited an MEI value significantly greater than zero (MEI = 1.09 ± 0.05; p < 0.001, cf. 0; Fig. 1c), reflecting an affinity for the membrane. Mutating the glycine residue at position 2 to alanine (PKA-Cn-G2A-mVenus), which prevented myristoylation (Clegg et al., 1989; Kamps et al., 1985; Towler et al., 1988), nearly completely abolished the membrane affinity (MEI = 0.08 ± 0.02, p < 0.001 cf. PKA-Cn-mVenus) (Fig. 1c). To evaluate the degree of membrane affinity of PKA-Cn-mVenus, its MEI was compared with that of mEGFP tagged KRas (Hancock et al., 1989), a known peripheral membrane protein. The MEI of PKA-Cn-mVenus was ∼45% of that of KRas (Fig. 1d), indicating that its membrane affinity is moderate. To estimate the fraction of PKA-Cn on the membrane, simulation was performed based on experimentally measured parameters. The point spread function (PSF) of our two-photon microscope was measured under the same conditions as the MEI experiments (20-50 µm deep in cultured hippocampal slices) (Figure 1e). The lateral and axial sizes of the PSF and the diameters of PKA-Cn-mVenus dendrites (ϕ = 2.18 ± 0.32; mean ± s.d.) were used to simulate how different fractions of membrane localization affected the image and MEI (Fig. 1f and Supplementary Fig. 1; see **Materials and Methods** for details). Using the measured MEI of PKA-Cn-mVenus on the working curve, ∼34% of the protein was estimated to be localized to the plasma membrane (Fig. 1f). This fraction on the membrane was likely underestimated, since PKA-Cn may also have affinity for intracellular membranes, which was not taken into account here. Overall, PKA-Cn was sufficient to confer a fractional membrane affinity in a myristoylation-dependent manner.

Because PKA-Cn alone was sufficient to confer association with the plasma membrane, if there was a second membrane-affinity motif, it must exist within PKA-Cn. Because PKA-Cn did not contain any additional lipid modification site, the presence of positively charged residues would be the only mechanism that could enhance the affinity of PKA-Cn to the typically negatively charged plasma membrane. There were eight basic residues (7 lysines and 1 arginine) and eight acidic residues (glutamate and aspartate) within PKA-Cn, resulting in an overall net neutral charge (Fig. 1a). Nevertheless, it is possible that the positive charges could not be counteracted by negative charges because of their unique positions and orientations in the 3D space. To address this possibility, we mutated the lysine residues to alanine residues two at a time and assayed their MEIs. Mutating the two lysine residues closest to the N terminus (K8,9A), but not those further away (K22, 24A and K29,30A, respectively), to alanine resulted in decreased MEIs (MEI = 0.63 ± 0.05 for PKA-Cn-K8,9A-mVenus, p< 0.001 cf. PKA-Cn-mVenus) (Fig 1b and 1c). Therefore, it appeared that only charges close to the N terminus modulated the membrane affinity of PKA-Cn. Indeed, another PKA-C N-terminal construct containing only the first 15 residues (named PKA-Cn15-mVenus) was localized to the membrane to a degree comparable to that of PKA-Cn (MEI = 1.19 ± 0.04 for PKA-Cn15-mVenus, p = 0.27, cf. PKA-Cn-mVenus) (Fig. 1c). The charge, rather than the residue identity, is likely the primary factor for the MEI decrease because when these lysine residues were mutated to similarly charged arginine residues (K8, 9R), the MEI was not altered significantly (Fig. 1d). Notably, even for PKA-Cn-K8,9A-mVenus, significant plasma membrane affinity remained (p < 0.001, cf. PKA-Cn-G2A-mVenus), suggesting that these residues quantitatively, rather than qualitatively, perturbed the membrane affinity of PKA-Cn.

The above results indicate that the two aforementioned lysine residues (K8 and K9) are the only candidates for the second membrane-affinity motif. However, within the first 15 residues of PKA-C, the net charge is also neutral, with these positively charged lysine residues are counterbalanced by two negatively charged glutamate residues (E12 and E14) (Fig. 1a). This net zero charge at the N terminus of PKA-C appears to be evolutionarily conserved from flies to mammals, and across different PKA-C isoforms (Fig. 1a, lower panel). We asked whether the two glutamates could effectively counter the effects of the K8 and K9 lysine residues with regard to plasma membrane affinity. If they could not, mutating these glutamates would have little or significantly smaller effects on the MEI compared to the lysine mutations. The opposite was found: mutating the glutamate residues to alanine or lysine (E12,14A and E12,14K, respectively) resulted in significantly increased MEIs (Fig. 1d). When combining the mutations at the lysine and glutamate positions, the membrane affinity is monotonically dependent on the summed charge from the four positions (Fig. 1d). The membrane fractions (Fig. 1g) and the Gibbs free energy relative to the wild-type PKA-Cn (Fig. 1h) were determined based on working curves computed using the corresponding dendrite sizes. A linear relationship was found between the charge and the binding energy regardless of whether the lysine or glutamate residues were mutated (R^2^ = 0.99, p < 0.001). Each positive charge contributed an energy of -0.89 kJ·mol^-1^. These results indicate that charges at the lysine and glutamate positions have equal weights in affecting the membrane affinity of PKA-Cn. Overall, although charges can modulate the membrane affinity, the observed affinity is intrinsic to myristoylation because PKA-Cn has a net zero charge.

We next examined whether full-length PKA-C is associated with the plasma membrane in a manner similar to that of PKA-Cn. PKA-C-mEGFP was co-expressed with untagged PKA-RIIα and the cytosolic marker mCherry. Consistent with our previous results using a different PKA regulatory subunit (PKA-RIIβ), full-length PKA-C was slightly excluded from the plasma membrane at rest as reflected by a negative MEI (MEI = -0.14 ± 0.03; p = 0.004, cf. 0) (Fig. 2a & 2b). This is likely because most PKA-C molecules at rest are bound to PKA-RIIα, the majority of which are in turn anchored to the microtubule cytoskeleton via MAP2 (Theurkauf and Vallee, 1982; Zhong et al., 2009). By increasing intracellular cAMP concentrations using the adenylyl cyclase activator forskolin along with the phosphodiesterase inhibitor IBMX, PKA-C translocated to become associated with the plasma membrane as reflected by a shift in the MEI to a positive value (MEI = 0.53 ± 0.03 upon stimulation; p< 0.001, both cf. rest and 0). In contrast, in separate experiments, mEGFP-tagged PKA-RIIα did not translocate under the same conditions (Fig. 2b), indicating that the fraction of PKA-C that translocated had dissociated from PKA-RIIα. This dissociation was consistent with the canonical understanding of PKA activation. The tendency to translocate to the plasma membrane upon stimulation was attenuated but not abolished when the K8 and K9 lysine residues were mutated to alanine (K8,9A), and were increased when E12 and E14 were mutated to lysines (E12,14K; Fig. 2b). The fraction of liberated PKA-C on the plasma membrane was estimated using an approach similar to that for the PKA-Cn mutants with two additional considerations: 1) PKA-C is slightly excluded from the membrane at rest (Fig. 2b), and 2) that not all PKA-C dissociated with the PKA-R under our stimulation conditions. To quantify the latter, we used two-photon fluorescent lifetime imaging microscopy (2pFLIM) (Ma et al., 2018; Yasuda et al., 2006) to measure the binding ratio between PKA-C-mEGFP and PKA-RIIα-sREACh (sREACh is a low-emitting yellow FP optimized for being a Foster resonance energy transfer acceptor to mEGFP for 2pFLIM) (Murakoshi et al., 2008). When normalized to the resting state, less than 70% (68 ± 5%) of PKA-C-mEGFP dissociated from PKA-RIIα-sREACh under our stimulation conditions (Fig. 2c). Using this dissociation fraction and the corresponding dendrite sizes (2.25 ± 0.39 µm for PKA-C-mEGFP, mean ± s.d.), ∼24.5% of liberated PKA-C was estimated to become associated with the plasma membrane. Since the membrane affinity of full-length PKA-C was not higher than that of PKA-Cn, it is unlikely that there is an additional membrane-affinity motif present in the remainder of PKA-C.

**Figure 2.**
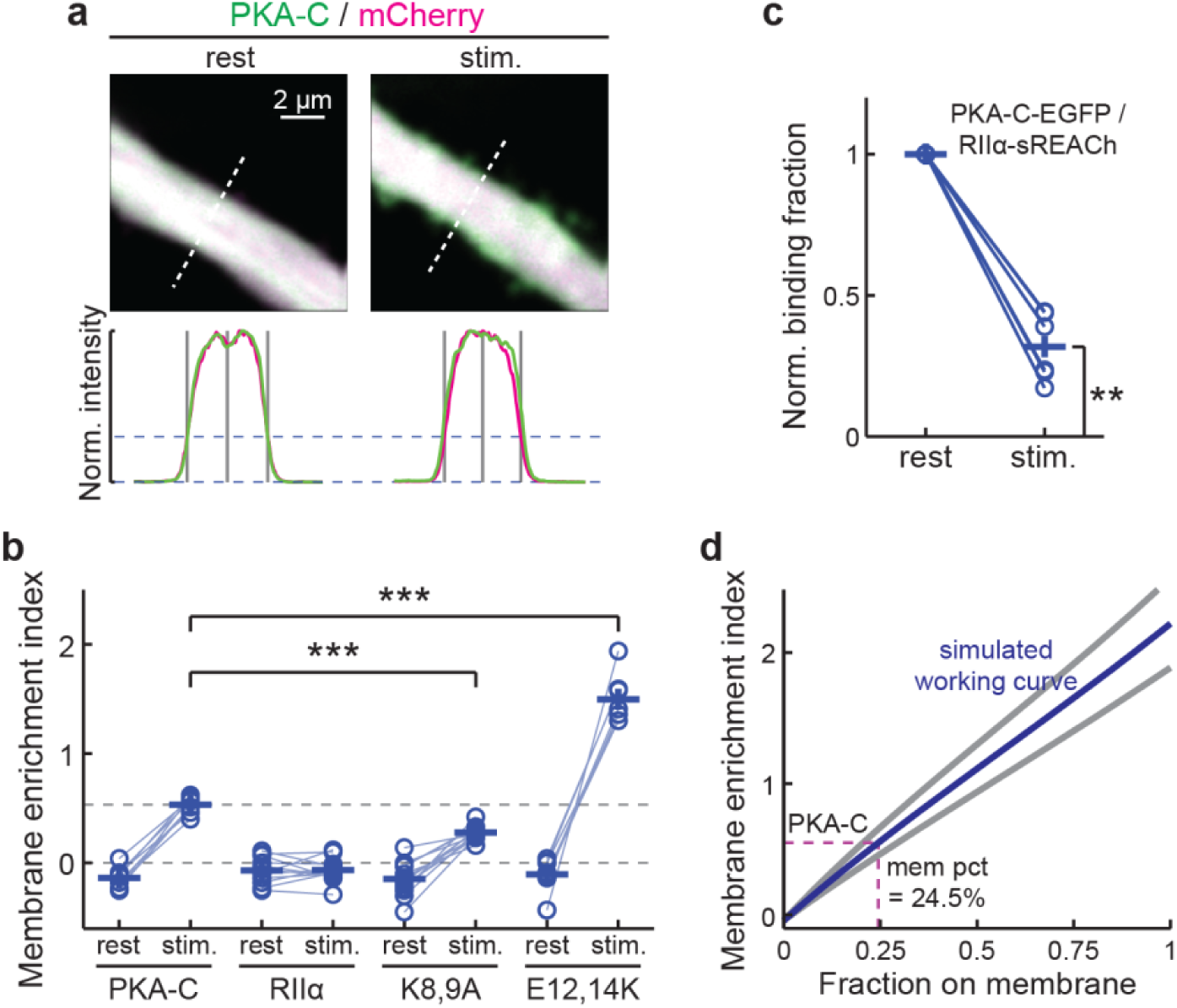
Association of full-length PKA-C with the plasma membrane. **a**, Representative images of thick neuronal apical dendrites and traces along the dashed white lines for quantifying MEIs for wild-type full-length PKA-C before (rest) and after stimulation (stim.) of PKA activity by forskolin (25 µM) and IBMX (50 µM). **b**, Collective MEI quantifications for the indicated full-length PKA-C constructs (co-expressed with untagged PKA-RIIα) and PKA-RIIα-mEGFP before and after forskolin and IBMX stimulation. From left to right, n = 8, 12, 12, and 7 for the indicated constructs. **c**, The binding ratio normalized to the resting state of PKA-C-mEGFP with RIIα-sREACh, as measured and quantified using 2pFLIM. **d**, MEI versus membrane fraction working curves using the experimental dendritic widths of PKA-C and the degree of PKA-C liberation, as measured in panel c. The estimated membrane fraction of liberated PKA-C is indicated (dash lines).

We used a second approach to confirm the tendency of PKA-C and its N-terminal mutants to associate with the membrane. Given that peripheral membrane-associated proteins diffuse much more slowly than cytosolic proteins (Harvey et al., 2008; Tillo et al., 2017), we assayed the protein motility of PKA-C, PKA-Cn, and their respective mutants. PKA-Cn and its mutants were fused to monomeric photoactivatable GFP (mPAGFP) (Patterson and Lippincott-Schwartz, 2002) and were co-expressed with the cytosolic marker mCherry or mCherry2. The mobility of these constructs was assayed by using focal two-photon photoactivation of the mPAGFP at individual spines and monitoring the resulting fluorescence decay over time (Fig. 3a and 3b) (Bloodgood and Sabatini, 2005; Gray et al., 2006). We focused on the spine because the time constant of the fluorescence decay at the spine (i.e., the spine-residence time) is linearly inversely proportional to the diffusion coefficient of a protein (Bloodgood and Sabatini, 2005). The G2A mutant of PKA-Cn exhibited a spine-residence time (τ = 0.30 ± 0.05) (Fig. 3c) comparable to previously reported cytosolic protein diffusion (Bloodgood and Sabatini, 2005; Tillo et al., 2017). The mobility of PKA-Cn was much slower (τ = 2.00 ± 0.15 s, p < 0.001, cf. G2A mutant) (Fig. 3c), which is consistent with its association with the membrane. The mobility of PKA-Cn was further decreased by the E12,14K mutations, which is consistent with a higher membrane affinity. While the mobility of PKA-Cn was increased by the K8,9A mutations, it was still much lower than that of the G2A mutant (Fig. 3c), corroborating the earlier conclusions based on MEIs that myristoylation is sufficient for membrane affinity in the absence of positively charged residues. Similarly, mutating K8 and K9 in full-length PKA-C increased its mobility but the increase was not to the same degree as that observed for the G2A mutants (Fig. 3d and 3e). These results supported the conclusions based on MEI measurements that the N-terminal fragment of PKA alone is sufficient for membrane association, and that the moderate membrane affinity can be modulated bi-directionally by changing charges near the myristoylation site.

**Figure 3.**
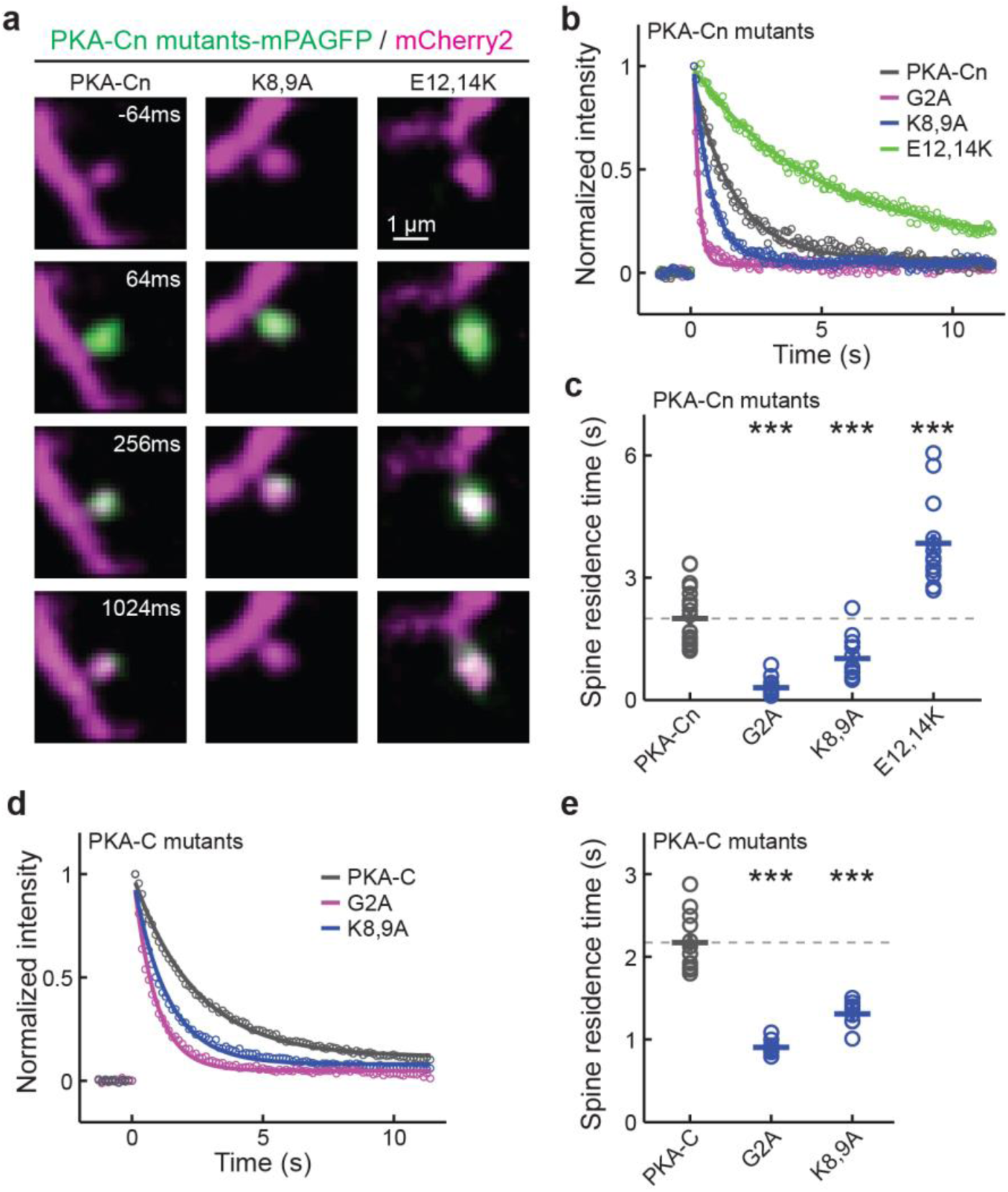
Mobility measurements of PKA-Cn and full-length PKA-C constructs. **a**, Representative time-lapse images of the indicated constructs. Two-photon focal activation of mPAGFP was carried out at time zero on a single spine. **b** & **c**, Representative traces (panel b) and collective results (panel c) of the spine residence time of the indicated PKA-Cn constructs. From left to right, n = 21, 15, 12, and 13. **d** & **e**, Representative traces (panel d) and collective results (panel e) of the spine residence time of indicated full-length PKA-C constructs co-expressed with untagged PKA-RIIα. Experiments were done in the presence of forskolin and IBMX to allow PKA-Cs to be liberated from PKA-Rs and translocate to the spine and the membrane. From left to right, n = 12, 9, and 8.

If only approximately one-fourth of liberated PKA-C is associated with the plasma membrane (Fig. 2d), can it explain the previous finding that PKA-C is more efficient in phosphorylating the same substrate on the membrane than in the cytosol (Tillo et al., 2017)? To address this question, we performed a computational analysis. Assuming that membrane-bound PKA-C can phosphorylate substrates up to 10 nm away from the membrane (∼size of PKA-C) (Zheng et al., 1993), and distributing the remaining liberated PKA-C evenly throughout the dendritic cytosol, with a dendrite size of 2.25 µm (the dendrite size used for Fig. 2d) and a membrane fraction of 24.5%, the PKA-C concentration at the membrane is more than 19-folds as high as its concentration in the cytosol (Fig. 4a). This explains the results of previous biochemical experiments that a PKA substrate is much more efficiently phosphorylated on the membrane than in the cytosol (Tillo et al., 2017).Overall, this analysis indicates that moderate membrane affinity of PKA effectively concentrates its protein activity at the plasma membrane.

**Figure 4.**
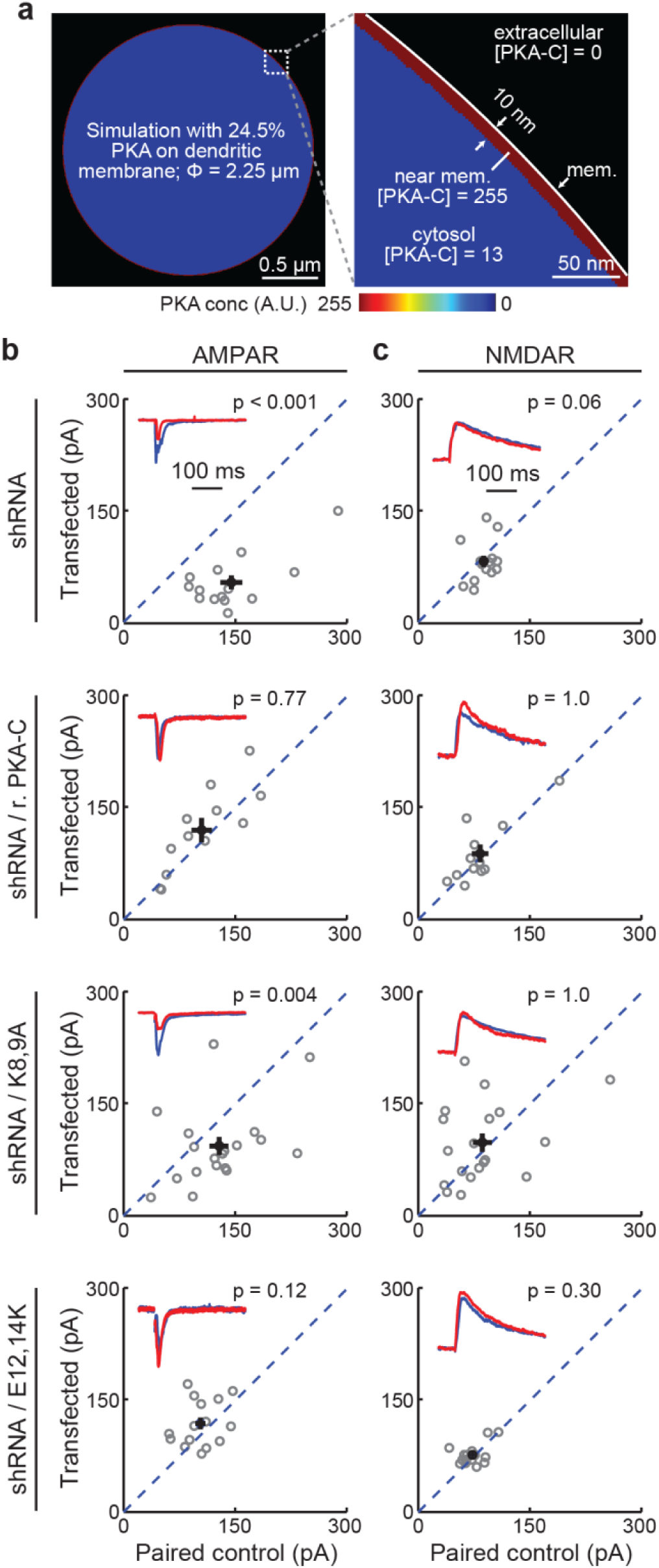
Fractional membrane association of PKA-C is important for its physiological function. **a**, Simulation experiment showing that the observed moderate membrane-affinity of full-length PKA-C is sufficient to enrich its activity at the membrane to over 19 times higher than its activity in the cytosol. **b** & **c**, Representative traces (red) normalized to the paired control (blue) (insets) and scatter plots of paired AMPA (panel b) and NMDA (panel c) receptor currents from neighboring untransfected CA1 neurons paired with those transfected with shRNA against PKA-C and the indicated shRNA-resistant rescue constructs. Statistical p values were obtained using a sign test (MATLAB). From top to bottom, n = 12, 14, 19, and 15.

Finally, is the moderate membrane affinity of PKA-C important for its regulation of synaptic functions? To address this, a previously established shRNA construct was used to selectively knock down PKA-Cα (Tillo et al., 2017). As shown previously, neurons expressing the shRNA construct exhibited significantly lower AMPA receptor (AMPAR) currents compared to paired, adjacent untransfected neurons (Fig. 4b1). NMDA receptor (NMDAR) currents were not statistically significantly altered under the same conditions (Fig. 4c1). As a result, the AMPAR/NMDAR current ratio was also reduced (Supplementary Fig. 2). The reduced AMPAR currents and reduced AMAPR/NMDAR current ratios were rescued by co-expression of an shRNA-resistant PKA-C-EGFP construct (r.PKA-C) (Fig. 4b and 4c and Supplementary Fig. 2). However, with decreased, but not abolished, membrane affinity, PKA-C bearing the K8,9A mutations failed to rescue the phenotype. In contrast, the E12,14K mutant, which exhibited an enhanced membrane-affinity, also rescued the phenotype. In addition, we have previously shown that PKA-C bearing the G2A mutation also cannot rescue knockdown of wildtype PKA-C in neuronal dendrites(Tillo et al., 2017). Taken together, these results indicated that, the membrane affinity resulting from myristoylation alone, while moderate, is important for normal synaptic function.

## Discussion

In summary, we found that the PKA-C N-terminal fragment is sufficient to associate liberated PKA-C, as well as functionally irrelevant FPs, to the plasma membrane in living neuronal dendrites. Because the PKA-C N-terminal fragment is both electrically neutral and lacks other membrane-affinity motifs, we concluded that myristoylation alone is sufficient to associate a protein to the plasma membrane in neuronal dendrites. Further, while the myristoylation only confers a moderate membrane affinity and only associates a fraction of PKA-C molecules with the plasma membrane, this fraction is sufficient to enrich PKA activity at the membrane by over an order of magnitude and is indispensable for physiological function. Taken together, these results suggest that the previous thought that myristoylation can only support functional membrane association in the presence of a second membrane-affinity motif must be revised. The revised view has broad implications in terms of understanding the wide range of proteins that are myristoylated (Farazi et al., 2001; Resh, 2013; Udenwobele et al., 2017; Wright et al., 2010).

Historically, PKA has not been considered to be a membrane-associated protein in part because: 1) it lacks a second membrane-affinity motif, which is traditionally deemed necessary for membrane association, 2) PKA-C appears to be highly soluble, and 3) the myristoylation of PKA-C does not appear to be essential for its catalytic function. Although this view was modified recently (Gaffarogullari et al., 2011; Tillo et al., 2017; Zhang et al., 2015), the above results remained to be reconciled. Here, we show that the membrane association PKA-C does not require a second membrane-affinity motif. At the same time, the membrane affinity of PKA-C is moderate, which explains its solubility. In addition, while the PKA catalytic function does not depend on the myristoylation, our results, together with previous results (Tillo et al., 2017) show that the fractional membrane affinity is still required for its successful regulation of synaptic function, presumably by concentrating its activity towards membrane substrates. It is also worth noting that small cellular compartments, such as neuronal dendrites, have a high membrane-to-cytosol ratio, which may have contributed to the detection of PKA-C at the membrane. Interestingly, although the membrane affinity of PKA-C appears to be tunable (e.g., by changing the negatively charged glutamate residues near the N terminus to alanine or lysine residues), its net zero charge is evolutionally conserved (Fig. 1a). Moderate membrane affinity may have advantages, such as maintaining certain levels of kinase activity in the cytosol, allowing the kinase to travel between discontinuous membranes, and allowing rapid inactivation of the kinase by regulatory subunits that are anchored away from the membrane.

## Materials and Methods

### Plasmid constructs

Constructs were made using standard mutagenesis and subcloning methods. All previously unpublished constructs and their sequences will be deposited to Addgene upon acceptance of the manuscript.

### Organotypic hippocampal slice cultures and transfections

Cultured rat hippocampal slices were prepared from P6 – P8 (typically P7) pups, as described previously (Stoppini et al., 1991; Zhong et al., 2009). Animal experiments were performed in accordance with the Guide for the Care and Use of Laboratory Animals of the National Institutes of Health, and were approved by the Institutional Animal Care and Use Committee (IACUC) of the Oregon Health & Science University (#IS00002274). cDNA constructs were transfected after 1.5–3 weeks *in vitro* via the biolistic gene transfer method using the Helios gene gun and 1.6 μm gold beads, or with single-cell electroporation (electroporation was used for Fig. 4 and Supplementary Fig. 2)(Otmakhov and Lisman, 2012), where long-term expression (∼ 1 week) was required.

### Two-photon imaging and photoactivation

A custom built two-photon microscope based on an Olympus BW51WI microscope body was used. Laser beams from two different Ti:Sapphire lasers (Maitai, Newport) were aligned to allow for simultaneous two-photon excitation and photoactivation. Laser intensities were controlled by Pockels cells (Conoptics). Imaging and photoactivation were controlled by ScanImage (Vidrio Tech) (Pologruto et al., 2003). Slices were perfused in gassed artificial cerebral spinal fluid (ACSF) containing 4 mM Ca, 4 mM Mg, and 0.5 µM tetrodotoxin (TTX) during imaging. MEI imaging experiments were carried out at 960 nm at the center z position of thick (> 1.5 µm, mean ∼2.2 µm) apical dendrites of CA1 neurons. mEGFP and mVenus fluorescence (green) were unmixed with that of the cytosolic marker (mCherry or mCherry2) using a dichroic (Chroma 565DCXR) and band-pass filters (Chroma HQ510/70 for green and Semrock FF01-630/92 for red). For in-tissue PSF measurements, 0.1-µm green fluorescent beads were puffed into tissue using a Picospritzer.

Photoactivation experiments were carried out using an 810 nm laser for focal photoactivation of mPAGFP and a separate imaging laser set at 990 nm to simultaneously image the activated mPAGFP (green) and the cytosolic marker (red). Spines expressing PKA-Cn constructs were imaged at 16 Hz at a 32×32 image size, and full-length constructs were imaged at 8 Hz with a 64×64 image size. Power for photoactivation was determined empirically at the time of individual experiments.

### Image analyses

Image analysis was performed using custom software written in MATLAB. For MEI measurements, line profiles of five-pixel width were manually drawn across the apical dendrites that were smooth on both sides (i.e., void of spines). The profiles were obtained from a single optical section centered along the middle z-plane of the dendrite. The background subtracted fluorescence intensity in the green (PKA constructs) and red (cytosol marker) channels were then used to calculate the MEI using the equation described in the main text.

Photoactivation experiments were analyzed by manually drawing ROIs over the x – y projection of the spines of interest. The green fluorescence emission from the ROI was then averaged and determined over each image frame of the experiment. Background values calculated from a manually drawn background ROI at the same frames were subtracted from the spine ROI values. Only spines that showed an activation signaling more than three times of the standard deviation of the baseline fluorescence fluctuation (before photoactivation) were included in data analysis. The green fluorescence intensity decays post-photoactivation were then averaged across three separate trials and subsequently fit with a single exponential decay. The spine residence time was defined as the time constant of the exponential fit.

### Computation and simulation

Computation and simulation were carried out in MATLAB. Simulation was used to generated the membrane fraction and MEI working curve. The full-width-half-maximum (FWHM) of averaged experimental PSFs was used to generate a Gaussian PSF with 10 nm pixel sizes in the x-z dimensions. The cytosolic distribution in the x-z dimensions was simulated as an evenly distributed round disk, and membrane distribution was simulated as signals being localized only at the edge of the disk. The diameter of the dendrite was specified by experimental measurements. Different percentages of signals were distributed to the membrane and the cytosol, and the resulting spatial distribution was convoluted with the PSF to produce the simulated images. MEI was measured at the z-center of the simulated image using the same algorithms as for experiments. For different constructs in Fig. 1g, the membrane fraction estimations were based on the corresponding average dendrite sizes. Gibbs free energy (Fig. 1h) was further calculated as:

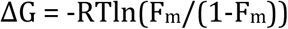

where F_m_ is the fraction on membrane, R is the gas constant, and T is the absolute temperature.

For Fig. 3d, the above simulation was modified to include one additional component of full-length PKA distribution: a fraction (31.8% of total) of immobile PKA-C was restricted from a zone that was 20 nm to the membrane. The restricted distribution by itself mimics the observed slightly negative MEI of PKA-C at rest. The fraction of mobile PKA-C on the membrane (with all mobile PKA-C summed to the 100%) is then plotted against MEIs to yield Fig. 3d.

For Fig. 4a, the pixel density was set to 2 nm and the membrane PKA activity was evenly distributed between *r* and *r*-0.01 µm of the circle (assuming that membrane-bound PKA-C can phosphorylate up to 10 nm [i.e., ∼ 1 PKA-C molecular size] away from the membrane), where *r* is the radius of the dendrite and was specified using the experimental data for PKA-C MEI measurements. The membrane distribution was then added to the cytosolic distribution where the remaining fraction of PKA-C is evenly distributed throughout the entire dendrite. The activity ratio between the membrane and cytosolic PKA concentrations can also be analytically derived as:

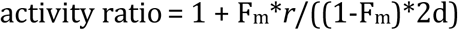

where F_m_ is the fraction of PKA on the membrane and d is the effective depth of membrane PKA phosphorylation. Computation and simulation gave essentially identical results.

### Two-photon fluorescence lifetime imaging and analyses

Two-photon fluorescence lifetime imaging microscopy (2pFLIM) was carried out in the time domain as described recently (Ma et al., 2018; Yasuda et al., 2006). Briefly, a two-photon microscope was modified by adding hardware components to compare the timing of each photon with the pulse timing of the Ti:Sapphire laser. Details of the setup will be provided upon request. Data acquisition was controlled by custom software (called FLIMimage) written in MATLAB provided by Dr. Ryohei Yasuda with modifications. Excitation and imaging conditions are similar to conventional two-photon imaging for MEIs with the exception that a Chroma ET500/40x barrier filter was used for green to allow better suppression of residual sREACh fluorescence in the green channel. Data analyses were performed using custom software written in MATLAB named FLIMview(Ma et al., 2018). Binding ratios were the result of fitting the fluorescence lifetime curve in FLIMview with two exponentials, with one reflecting the unbound state and the other reflecting the bound state. FLIMview will be made available upon reasonable requests.

### Electrophysiology

Whole-cell voltage-clamp recordings were performed using a MultiClamp 700B amplifier (Molecular Devices). Electrophysiological signals were filtered at 2 kHz and digitized and acquired at 20 kHz using custom software written in MATLAB. Slices were perfused with artificial cerebrospinal fluid containing 4 mM Ca and 4 mM Mg. The internal solution contained (in mM) 132 Cs-gluconate, 10 HEPES, 10 Na-phosphocreatine, 4 MgCl2, 4 Na2-ATP, 0.4 Na-GTP, 3 Na-ascorbate, 3 QX314, and 0.2 EGTA with an osmolarity of 295 mOsmol/kg. The junction potential was calculated to be -17 mV using a built-in function in the Clampfit software (Molecular Devices). Several less abundant anions (phosphocreatine, ATP, GTP and ascorbate) were omitted in the calculation due to lack of data in the program. The Cl reversal potential was -75 mV.

To reduce recurrent activities, cultured hippocampal slices were cut on both sides of CA1 and 4 µM 2-chloroadenosine (Sigma) was present in all recording experiments. 10 µM GABAzine (SR 95531, Tocris) was also included to suppress GABA currents. For electrical stimulation, a bipolar, θ-glass stimulating electrode (Warner Instruments) was positioned in the stratum radiatum 100–150 μm lateral to the recorded neuron. For all recordings, a transfected neuron and an untransfected neuron located within 50 µm of each other were sequentially recorded without repositioning the stimulation electrode. Measurements were carried out on averaged traces from approximately 20 trials under each condition. For AMPAR currents, the cells were held at -60 mV (before correcting for the junction potential) and the current was measured as the baseline-subtracted peak current within a window of 2–50 ms after electric stimulation. activation. For NMDAR currents, the average currents at 140 to 160 ms after stimulation were used when the cells were held at +55 mV (before correcting for the junction potential)

### Data analysis, presentation, and statistics

Quantification and statistical tests were performed using custom software written in MATLAB. Averaged data are presented as mean ± s.e.m., unless noted otherwise. Throughout the paper, “n” indicates the number of neurons for MEI analysis, and the number of spines for photoactivation experiments, unless noted otherwise. For photoactivation experiments, no more than four spines were from a single neuron. p values were obtained from one-way ANOVA tests, unless noted otherwise. In all figures, *: p ≤ 0.05 and is statistically significant after Bonferroni correction for multiple tests, **: p ≤ 0.01, and ***: p ≤ 0.001.

## Acknowledgements

We thank all members of the Mao and Zhong laboratories at the Vollum Institute for constructive discussions. We thank Drs. Tianyi Mao, John Williams, Wolfhard Almers, and Bart Jongbloets for critical comments on the manuscript. This work was supported by two NIH BRAIN Initiative awards to H.Z. (U01NS094247 and R01NS104944).

## Author contributions

H.Z. conceived of the project. W.X. and H.Z. designed the experiments. W.X. and H.Z. performed the experiments and data analyses. M.Q. produced reagents used in the experiments. H.Z. wrote the manuscript with comments from W.X.

---

**Supplementary Figure 1.**
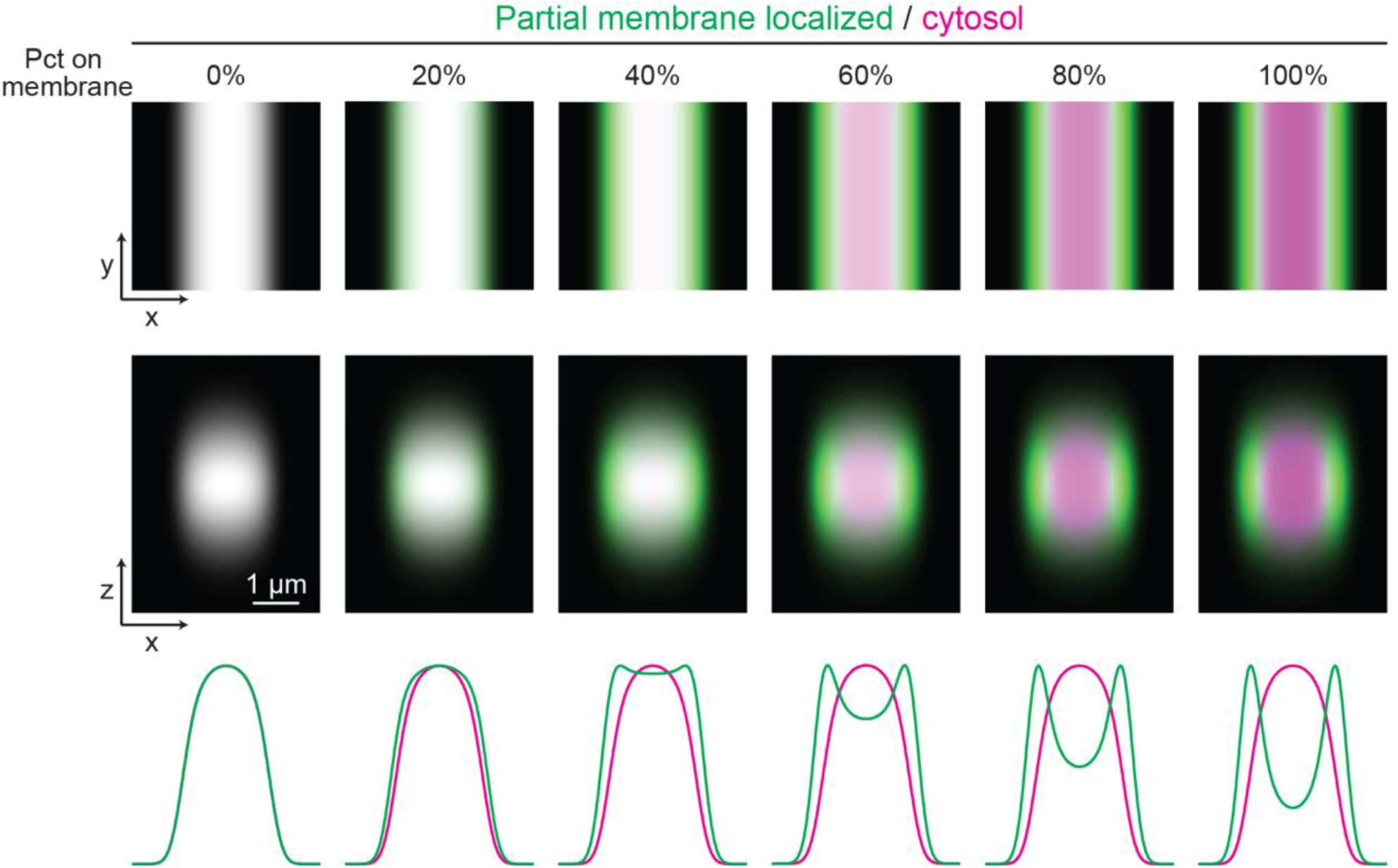
Representative simulated images and traces of protein localization with different fractions on the membrane. Proteins (green) with indicated percentage (pct) on the membrane of a model cylindrical neuronal dendrite (ϕ = 2.18 µm) along the y axis, with the remaining evenly distributed in the cytosol, are convoluted with a Gaussian simulated point spread function (PSF) with a lateral full-width-half-maximum (FWHM) size of 0.505 µm and an axial size of 1.78 µm. Both the x-y view (top) and the x-z view (middle) are shown and overlaid on the image of a fully cytosolic marker (magenta). Bottom panels show the normalized fluorescence intensity traces along the x axis at the z center.

**Supplementary Figure 2.**
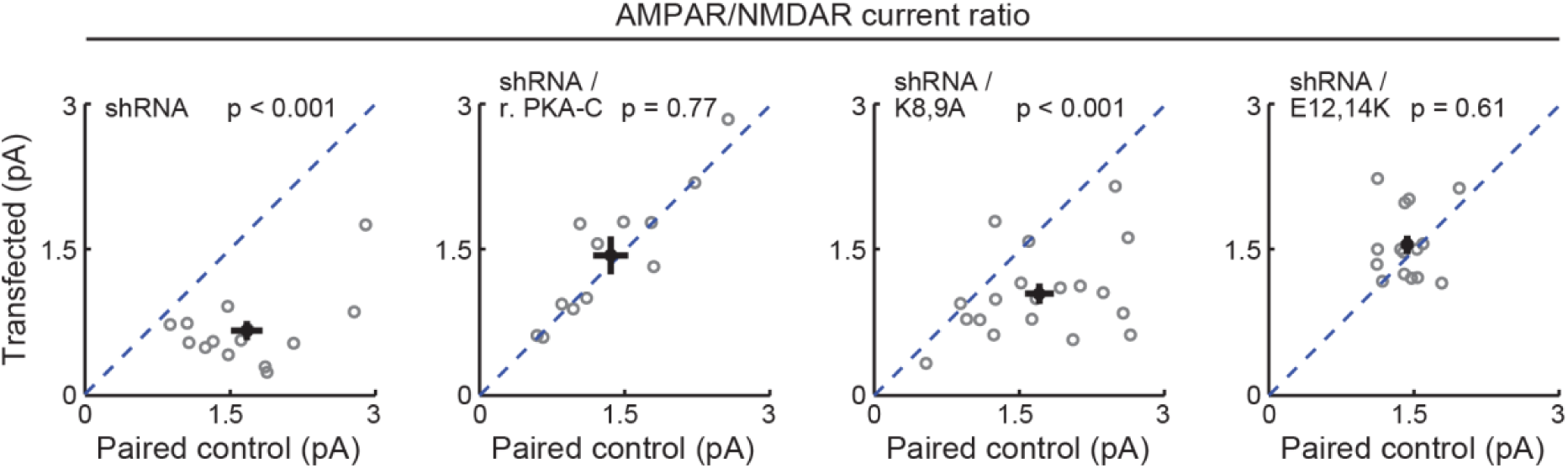
AMPA/NMDA receptor current ratio. Scatter plots of paired AMPA receptor to NMDA receptor current ratios from neighboring untransfected CA1 neurons paired with those transfected with shRNA against PKA-C and the indicated shRNA-resistant rescuing constructs. Statistical p values were tested using a sign test (MATLAB). From left to right, n = 12, 14, 19, and 15.

